# Formulation Development of Topical Inserts Containing Doxycycline and Doxycycline Combined with Tenofovir Alafenamide and Elvitegravir for the Prevention of Sexually Transmitted Infections

**DOI:** 10.64898/2026.02.06.704457

**Authors:** Vivek Agrahari, M. Melissa Peet, Jasmin Monpara, Rijo John, Sriramakamal Jonnalagadda, Pardeep K. Gupta, Meredith R. Clark, Gustavo F. Doncel

## Abstract

**Purpose:** Despite advances in oral and injectable HIV prevention options and oral prophylaxis for sexually transmitted infections (STIs) of bacterial origin, there remains a critical need for effective on-demand topical (vaginal/rectal) products for pre- and post-exposure prophylaxis (PrEP and PEP). To fill this gap, we have developed single and first-in-kind multi-active topical inserts for bacterial STIs and HIV/STIs prevention.

**Methods:** We have formulated two different inserts, one containing doxycycline (DOX) at 10, 50, and 100mg doses for bacterial STI prevention, and a multipurpose prevention product (TED insert) that combines DOX (10mg) with the antiretrovirals tenofovir alafenamide (TAF; 20mg) and elvitegravir (EVG; 16mg) to target both bacterial STIs and HIV.

**Results:** Inserts were manufactured through a simple, cost-effective process. Drug loading was within 95-105% of the labeled amount, confirming a robust manufacturing process. In vitro, they disintegrated within 10min with >95% drug release within 60min. The dissolution behavior of DOX inserts showed surface erosion but was affected by medium volume and drug amount. The inserts met key physicochemical targets: hardness (5-8kg), friability (<1%), moisture content (<2%), and osmolality (<550mOsm/kg). Based on 6-month storage stability, DOX inserts maintained their physicochemical properties, suggesting a shelf life of >2years. Preliminary 1-month stability of TED inserts under accelerated conditions showed preservation of their physicochemical properties.

**Conclusion:** This study represents the first formulation development report on topical inserts containing DOX alone or in combination with antiretrovirals. Both inserts offer a novel, on-demand topical STI prevention option that supports flexible PrEP/PEP use by both women and men.

## INTRODUCTION

Human immunodeficiency virus (HIV) and sexually transmitted infections (STIs) of bacterial origin, including syphilis, chlamydia, and gonorrhea, remain major global public health challenges. In 2023, an estimated 39 million people worldwide were living with HIV, with about 1.3 million new infections occurring annually [1]. Bacterial STIs are even more common, and the World Health Organization (WHO) estimates over 374 million new cases of curable STIs (syphilis, chlamydia, gonorrhea, trichomoniasis) occur each year [2]. These epidemics are interlinked; the presence of bacterial STIs can increase both susceptibility to and transmissibility of HIV through mechanisms such as mucosal disruption, inflammation, and increased viral shedding [3–7]. While consistent condom use remains the most broadly protective prevention tool against both HIV and bacterial STIs, additional biomedical strategies have emerged. For HIV, these include oral and injectable pre-exposure prophylaxis (PrEP), treatment as prevention (TasP), and post-exposure prophylaxis (PEP) [8–11]. For bacterial STIs, biomedical preventatives are limited. However, doxycycline (DOX) oral PEP has recently shown efficacy in reducing chlamydia and syphilis in cisgender men and transgender women when taken near the time of sexual exposure [12–14]. When the same oral regimen was tested for efficacy in cisgender women, the incidence of STIs was not significantly lower; however, hair sample drug analysis confirmed that adherence was low [15]. These findings have nonetheless influenced the US public health agency’s proposed guidelines for the use of DOX as PEP taken within 72 h following possible STI exposure at an oral dose of 200 mg [16]. Together, these data highlight both the progress and remaining gaps, especially for desirable products for cisgender women, in integrated strategies to reduce HIV and STI burdens. One clear gap is the lack of on-demand, topical options that may prevent infection, offering user-control, event-driven protection, limited drug systemic exposure, and reduced safety concerns, overall representing an appealing method to certain populations, for instance, those having infrequent or intermittent sex. Surveys conducted in sub-Saharan Africa by the MATRIX consortium show support for this topical on-demand option among adolescent girls and young women (unpublished data). Published studies have demonstrated that many women value flexible, on-demand STI and HIV prevention options that align with episodic sexual activity, underscoring the importance of user-controlled, event-driven prevention strategies [17,18].

To this end, CONRAD embarked on the development of an on-demand dual-compartment insert containing tenofovir alafenamide fumarate (TAF) and elvitegravir (EVG) for HIV prevention [19,20]. The TAF/EVG (20 mg/16 mg) insert was successfully scaled up from laboratory development to clinical manufacturing under current Good Manufacturing Practices (cGMP) conditions. The formulation demonstrated preclinical efficacy against HIV-1 and successful Phase I clinical trials [20–22]. Leveraging these results and the drug delivery platform, we further sought to develop an on-demand insert containing DOX alone and combined with TAF/EVG for the prevention of bacterial STIs and HIV/STIs (multipurpose prevention technology; MPT), respectively. Similarly to the clinically tested TAF/EVG inserts, the DOX and TAF/EVG/DOX (TED) inserts described here are solid dosage forms suitable for retention in a body cavity, for instance, following vaginal and/or rectal applications, and are therefore categorized as “topical inserts” for their dual-compartment applicability, as discussed in our previous publication [23].

This manuscript focuses on the development, physicochemical characterization, and stability evaluation of DOX vaginal inserts formulated at various strengths (10, 50, and 100 mg) for direct application to the main site of primary infection, aiming to prevent STIs before or after exposure. Based on the DOX oral dosage and pharmacokinetics (PK) studies [24–26], we selected three doses of DOX as a starting point to test preclinically with the goal of exceeding the in vitro MIC_90_ range for *Chlamydia trachomatis* (0.016-0.064 µg/mL) [27], *Neisseria gonorrhoeae* (16 μg/mL) [28], and *Treponema pallidum* (0.06-0.10 µg/mL) [29] in mucosal fluids to ensure efficacy. We also evaluated the combination of TAF, EVG, and DOX in a single insert formulation (TED insert) and analyzed their physicochemical characteristics and pilot stability to assess the formulation development feasibility. The co-formulation of two synergistic but mechanistically distinct ARVs (TAF and EVG) with DOX into a single product enables extended STI/HIV protection from a single, before or after sex, application. DOX and TED inserts met the evaluation criteria for manufacturability and in vitro characterization and demonstrated several promising attributes, including a simple and robust direct-compression manufacturing process, rapid initial disintegration and drug dissolution, adequate mechanical strength, and long-term stability. Visual assessment of DOX inserts showed dissolution via surface erosion, enabling consistent drug release; however, dissolution behavior depended on the dissolution medium volume and drug load.

To our knowledge, this study is the first report describing the development of a topical (vaginal/rectal) insert containing either DOX alone or in combination with ARVs. Furthermore, no topical insert platform has yet been reported that incorporates three distinct drugs to address multiple STI indications, underscoring the novelty of the TED inserts described in this paper. Our initial formulation development efforts described herein have focused on the vaginal route, with future efforts targeted to further cover the rectal route of administration. Proof-of-concept testing in non-human primates (NHPs) is ongoing in collaboration with the Centers for Disease Control and Prevention (CDC) to assess PK and efficacy [30]. The robust formulation development and encouraging in vitro performance of the DOX and TED inserts, together with successful clinical translation and promising Phase I trial outcomes of our prior TAF/EVG inserts, support the potential of these DOX and TED inserts as safe, efficacious, and cost-effective on-demand alternatives to existing STI and HIV prevention strategies, addressing a critical unmet need.

## MATERIALS AND METHODS

### Materials

Doxycycline hyclate (DH) was purchased from Medisca Inc. (Plattsburgh, NY), and tenofovir alafenamide fumarate (TAF) was purchased from VulcanChem (Pasadena, CA). Elvitegravir (EVG) was kindly provided by Gilead Sciences, Inc. (Foster City, CA) through material transfer agreements with CONRAD/Eastern Virginia Medical School. Povidone (K-29/32) was obtained from Ashland (New Jersey, NJ), while lactose anhydrous, mannitol, and magnesium stearate were purchased from Spectrum Chemical (Brunswick, NJ). Poloxamer 188 was acquired from Sigma-Aldrich (St. Louis, MO), and poly(ethylene glycol) (PEG) 8000 from Fisher Scientific (Hampton, NH). All other chemicals were of high-purity analytical grade and used as is.

### High-Performance Liquid Chromatography (HPLC) Analysis

#### DOX inserts

The HPLC system (Agilent Technologies, Santa Clara, CA) included a binary pump, a temperature-controlled autosampler, a YMC-Pack Pro C18 column (150 × 4.6 mm ID, 3 µm particle size) (YMC America, Allentown, PA), and a UV detector. An isocratic elution with mobile phase A (100% v/v acetonitrile) and mobile phase B (0.25% v/v perchloric acid, pH 2.5, adjusted with 5 M sodium hydroxide) at a 26:74 % v/v ratio was used, with a flow rate of 1 mL/min. Each sample was injected at 20 µL, with a runtime of 30 min and detection at 350 nm. Data were acquired and processed using ChemStation rev. B.03.02 LC software (Agilent Tech.). The standard stock solution of DOX (DH) at 1 mg/mL was prepared in a diluent (26:74% v/v ratio of mobile phase A to B). Calibration standards in the concentration range of 7.8-500 µg/mL were prepared by serial dilution of this stock solution using the same diluent to test linearity.

#### TED inserts

The Agilent HPLC system was equipped with an Agilent Zorbax C18 column (150 × 4.6 mm ID, 3 µm particle size). Gradient elution was used at a flow rate of 1 mL/min at ambient temperature. The HPLC elution time program with the mobile phase A (100% v/v acetonitrile) and mobile phase B [(10mM potassium hydrogen phosphate, pH 2.5, adjusted with orthophosphoric acid (OPA: 85% *v*/v)] was as follows: 0.01 min, 95% B and 5% A; 6 min, 20% B and 80% A; 8 min, 20% B and 80% A; 9 min, 95% B and 5% A; and 10 min, 95% B and 5% A. Each sample was injected at 5 µL, with a runtime of 10 min and detection at 255 nm. Standard stock solutions of the drugs were prepared at 0.5 mg/mL in a 90:10 (v/v) methanol:water diluent. Serial dilutions of each drug’s stock solution in the same dissolution medium yielded calibration and working standards over the concentration range of 3.1-225 µg/mL.

For both DOX and TED inserts, the HPLC autosampler and column were maintained at 4°C and 25°C, respectively. System suitability was confirmed through repeatability testing (drug peak area response and retention time), assessment of drug peak interference with the diluent, and drug recovery testing, based on percent relative standard deviation (% RSD).

### Inserts Formulation Development

#### DOX inserts manufacturing and packaging

Three different DOX formulations containing DH equivalent to 10, 50, and 100 mg of DOX free base (D1, D2, and D3, respectively) were produced by direct compression using a rotary tablet press (MTP-4, Mendel, East Hanover, NJ). Briefly, the drug, along with excipients (except magnesium stearate), was screened through a #20 stainless steel sieve. The sieved material was transferred to a jar, blended for 5 min with rotation and tumbling, and then moved to the tablet machine hopper. The slug (1-inch diameter and 4 mm thick) was compressed to 1-2 kg hardness and then passed through a #16 screen sieve. The necessary amount of material was transferred into a jar, and magnesium stearate was added. This mixture was then blended, rotated, and tumbled for 2 min. The DOX inserts were compressed using this blend, while in-process checks for weight and hardness were performed. The inserts were packed in 30 cc high-density polyethylene (HDPE) bottles (MRP Solutions, Somerset, NJ), each containing 20 inserts with cotton and 1 g of silica gel desiccant (Wisesorbent Tech. LLC, Marlton, NJ). The HDPE bottles were induction-sealed with a GLF-500 electromagnetic induction foil capping machine using laminated plastic foil bottle cap liners (EEZglobal Inc., Hayward, CA).

#### TED inserts manufacturing and packaging

The inserts were manufactured using a similar direct-compression process to that of the DOX inserts, with the following modifications to accommodate three drugs. Briefly, the ingredients were passed through a #30 stainless steel sieve in the following order: 50% lactose, TAF, EVG, DOX, povidone, poloxamer, PEG, mannitol, and the remaining 50% lactose. The sieved material was transferred to a 2-qt V-shell blender and blended for 5 min at 25 rpm. Magnesium stearate (0.4% w/w of the total amount in the final batch) was accurately weighed, passed through a #30 mesh screen, and then added to the remaining blend, which was mixed for 3 min using the same blender. Using this blend, the slug (4 mm thick) was compressed to a hardness of 2-3 kg, then passed through a #20 sieve. The remaining magnesium stearate (1.1% w/w of the total amount) was weighed, passed through a #30 mesh screen, and added to the blend. This mixture was then blended, rotated, and tumbled for 3 min. The TED inserts were compressed using this blend. The inserts were packed in 30 cc HDPE bottles (induction-sealed), each containing 25 inserts with cotton and 3 g of silica gel desiccant.

### Physicochemical Characterization and Stability Testing of Inserts

#### Hardness and friability testing

To evaluate hardness, ten inserts (n = 10) were examined using an HT-300 hardness tester (Key International, Inc., Englishtown, NJ). Each insert was subjected to an increasing load applied along the radial axis until the insert broke or fractured. For friability assessment, thirteen inserts (n = 13) were weighed and loaded into a friabilator (Distek, Inc., North Brunswick, NJ), which was run at 25 rpm for 4 min. The inserts were dusted off and reweighed. The average percentage weight loss was determined by calculating the difference between the initial and final weights using the following equation. Here, I_w_ and F_w_ are the total initial and final weights of the inserts, respectively.

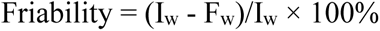

#### Moisture content analysis

Inserts (n = 3) were crushed separately into a fine powder using a mortar and pestle. A 50 mg equivalent of powder from each insert was dissolved in 5 mL of methanol and analyzed by Karl Fischer coulometric titration (Mettler-Toledo, Columbus, OH).

The water content was measured as a numerical value, expressed as the percentage of the sample’s weight before and after injection into the Titrator.

#### Osmolality Testing

An Advanced Micro Osmometer Model 3300 (Advanced Instruments, Inc., Norwood, MA) was used for measuring osmolality. The instrument was calibrated with a sodium chloride solution (Wescor, Inc., Logan, UT) as the standard. Each sample was added to SVF (pH ∼4.3, 5 mL) and sonicated for ∼60 min. A 20 µL aliquot was drawn and injected into the central depression of the osmometer holder, which was then placed into the instrument. The osmolality was recorded in mOsm/kg (n = 3).

### Drug content assay

DOX inserts (n = 10) were accurately weighed, crushed, and triturated to a fine powder using a mortar and pestle. A quantity of insert powder equivalent to 10 mg of DOX was weighed and added to 20 mL of diluent, a 26:74% v/v mixture of the HPLC mobile phases A and B. This mixture was sonicated for 30 min, and a 1 mL aliquot was centrifuged at 5,000 rpm for 10 min. The supernatant was collected and analyzed by HPLC.

For TED inserts, 10 inserts were crushed and triturated to a fine powder using a mortar and pestle. A powder equivalent to 20 mg of TAF free base, 16 mg of EVG, and 10 mg of DOX (free base) was weighed and added to 10 mL of water. This was sonicated for 5 min to disperse the powder, followed by the addition of 80 mL of methanol and an additional 10 min of sonication to dissolve the content. The final volume was adjusted to 100 mL with methanol, and the drug content was determined by HPLC.

### Disintegration and dissolution (drug release) testing

The disintegration analysis of inserts (n = 6) was performed using a DT 2V 115 disintegration tester (SOTAX GmbH, Lörrach, Germany). A 10-mesh screen (2 inches) was placed at the bottom of the beaker, and 1 L of water was added while maintaining a temperature of 37°C with a rotation speed of 30 rpm. The inserts were placed onto the mesh, and the time to complete disintegration (as visually observed) was recorded. For the dissolution and drug release testing of DOX inserts (n = 6), a paddle-type apparatus 2 (Distek, Inc., North Brunswick, NJ) was used in accordance with the United States Pharmacopeia (USP) method. The acetate buffer (900 mL) at pH ∼4.5 was used as the dissolution medium, mimicking the pH of cervicovaginal fluid. The dissolution apparatus, equipped with a paddle speed of 50 rpm, was operated at 37°C. Dissolution samples (1 mL) collected at various time points, up to 60 min, were measured for the cumulative release of DOX using BioTek Synergy 2 Microplate Reader (Agilent Techn., Santa Clara, CA) at 268 nm. A serial dilution of the stock solution with the same dissolution medium yielded calibration standards in the concentration range of 7.5-120 µg/mL for dissolution testing. The calibration curve was linear, with a correlation coefficient (R²) of 0.999.

To analyze the disintegration and dissolution of DOX inserts simulating in vivo conditions, inserts were exposed to 1 and 3 mL of simulated vaginal fluid (SVF, pH ∼4.3 at 37°C, mimicking the vaginal environment) prepared using a published protocol [31], and 4 mL of a simulant fluid mixture (pH ∼7.2 at 37°C, mimicking the environment during sexual intercourse) consisting of 1 mL of SVF mixed with 3 mL of simulated semen fluid (SFS, pH ∼7.8) [32]. Each insert was placed on a Petri dish, and the respective volumes of SVF or simulant mixture (SVF + SFS) were added on top of the insert. Images were captured at predetermined time intervals of 0, 5, 10, 15, 30, 45, 60, 90, and 120 min, using a polarized Microscope Digital Camera (MU1000, AmScope, China) to monitor the visual changes on the inserts.

The dissolution of TED inserts was also studied using a USP method similar to that for DOX inserts (900 mL of acetate buffer at pH 4.5), except that 0.5% w/v sodium lauryl sulfate (SLS) was added to the dissolution medium. One milliliter of the filtered sample was withdrawn at various timepoints, up to 120 min, and the drug concentrations were analyzed by HPLC.

#### Stability testing

The stability studies of DOX inserts were conducted for 6 months under controlled room conditions (25°C/60% RH) and accelerated conditions (40°C/75% RH) following the International Council for Harmonisation (ICH) guidelines Q1A(R2) [33]. Insert samples packaged in HDPE bottles with cotton and silica gel desiccant were collected at different time points and analyzed for drug content and physicochemical properties. To confirm proof of concept, the stability of TED inserts was evaluated for 1 month under accelerated conditions (40°C/75% RH) and compared with time-zero data. The inserts were analyzed for physical appearance, drug assay, hardness, friability, and drug dissolution.

### Statistical Analysis

GraphPad Prism version 10.3 was used to analyze and graph the data. Results were calculated as the mean value ± standard deviation (SD) unless specified. The number of replicates used in each study is mentioned in its respective Figure and Table.

## RESULTS

### DOX Inserts Formulation Development and Characterization

#### HPLC method development

The representative HPLC chromatogram of DOX (DH: retention time: RT ∼15.79 min) is shown in **Supplementary Fig. 1.** The calibration curve in the range of 7.8-500 µg/mL was linear with a correlation coefficient (R^2^) of 0.9995. For the method validation parameters (system suitability, repeatability, and drug recovery), the %RSD was <2%. No interference from the diluent peak was observed at the drug’s retention time or in its peak area. ***Insert manufacturing and physicochemical characterization:*** Three different formulations containing low (10 mg: D1), medium (50 mg: D2), and high (100 mg: D3) amounts of DOX **(Table 1)** were produced, each with a targeted 500 mg insert weight **(Fig. 1)**. For the excipients, mannitol, lactose, PEG, poloxamer, povidone, and magnesium stearate were rationally selected due to their favorable physicochemical properties, which facilitated insert manufacturing via direct compression.

**Fig. 1.**
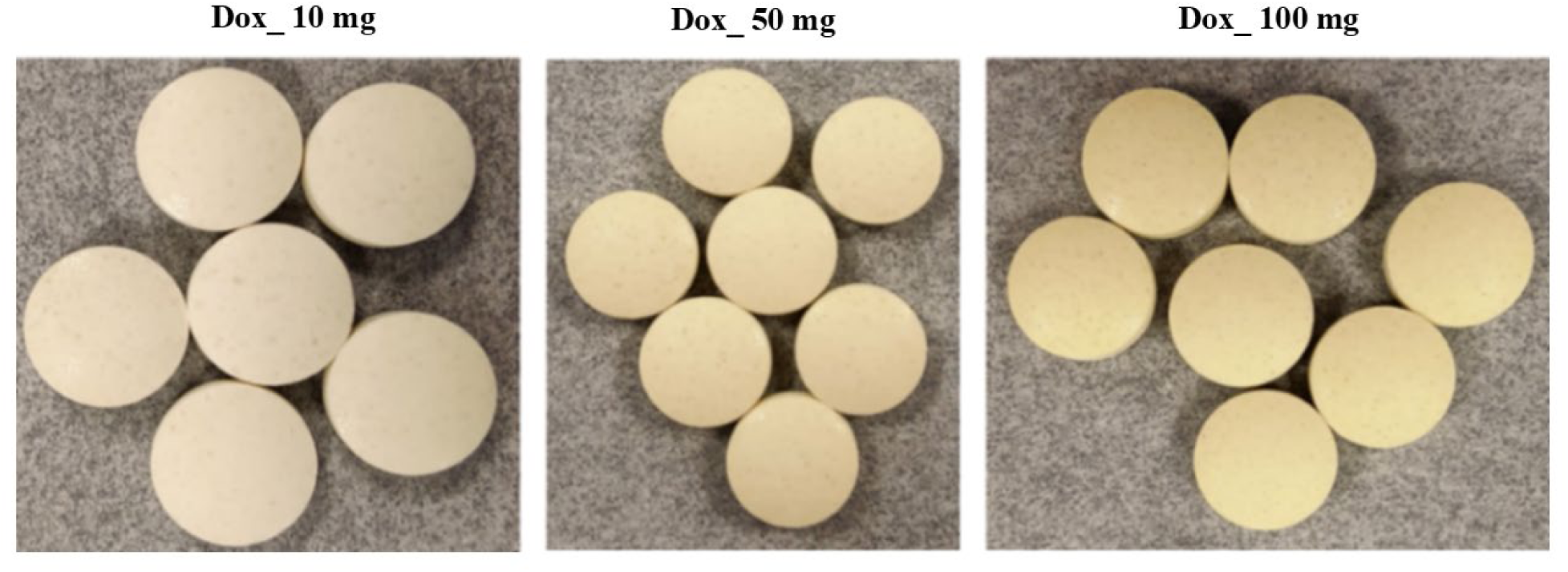
DOX inserts (500 mg) at 10, 50, and 100 mg strengths.

**Table 1:** Compositions of DOX inserts.

| Ingredients | Insert formulations |  |  |  |  |  |
| --- | --- | --- | --- | --- | --- | --- |
|  | D1 (10 mg DOX) |  | D2 (50 mg DOX) |  | D3 (100 mg DOX) |  |
|  | % w/w | Weight (mg) | % w/w | Weight (mg) | % w/w | Weight (mg) |
| DOX hyclate (DH) | 2.31 | 11.55* | 11.55 | 57.75* | 23.13 | 115.65* |
| Povidone (K-29/32) | 4.00 | 20.00 | 4.00 | 20.00 | 4.00 | 20.00 |
| Poloxamer 188 | 2.00 | 10.00 | 2.00 | 10.00 | 2.00 | 10.00 |
| Lactose anhydrous | 58.19 | 290.95 | 48.95 | 244.75 | 37.37 | 186.85 |
| Mannitol | 17.00 | 85.00 | 17.00 | 85.00 | 17.00 | 85.00 |
| PEG 8000 | 15.00 | 75.00 | 15.00 | 75.00 | 15.00 | 75.00 |
| Magnesium stearate | 1.50 | 7.50 | 1.50 | 7.50 | 1.50 | 7.50 |
| Total | 100 | 500 | 100 | 500 | 100 | 500 |

\* Equivalent to 10 mg (D1), 50 mg (D2), and 100 mg (D3) of DOX.

The physicochemical characteristics of DOX inserts are summarized in **Table 2**. The formulations showed acceptable results for drug assay (within 95-105% of the initial drug content), hardness (within 5-8 kg), friability (<1%), and moisture content (<2%). In 5 mL of SVF, DOX inserts at all three dose levels exhibited an osmolality below 550 mOsm/kg. Under the USP testing method, inserts began to disintegrate within 5 min, and complete disintegration was observed within ∼10 min, achieving nearly 100% DOX release within 60 min (over 85% within 30 min) of dissolution. Visual assessments **(Fig. 2)** showed that the dissolution of 10 mg DOX inserts in 3 mL media volume (SVF or simulant mixture) was predominantly governed by surface erosion **(Fig. 2B and 2C)**. However, when tested in 1 mL of SVF, the same insert demonstrated minimal erosion over the same period **(Fig. 2A)**. Similarly, the 50 mg DOX inserts exhibited limited surface erosion **(Fig. 2D-F)** when placed in 1 or 3 mL of SVF, with increased disintegration in 4 mL of mixed simulant fluids.

**Fig. 2.**
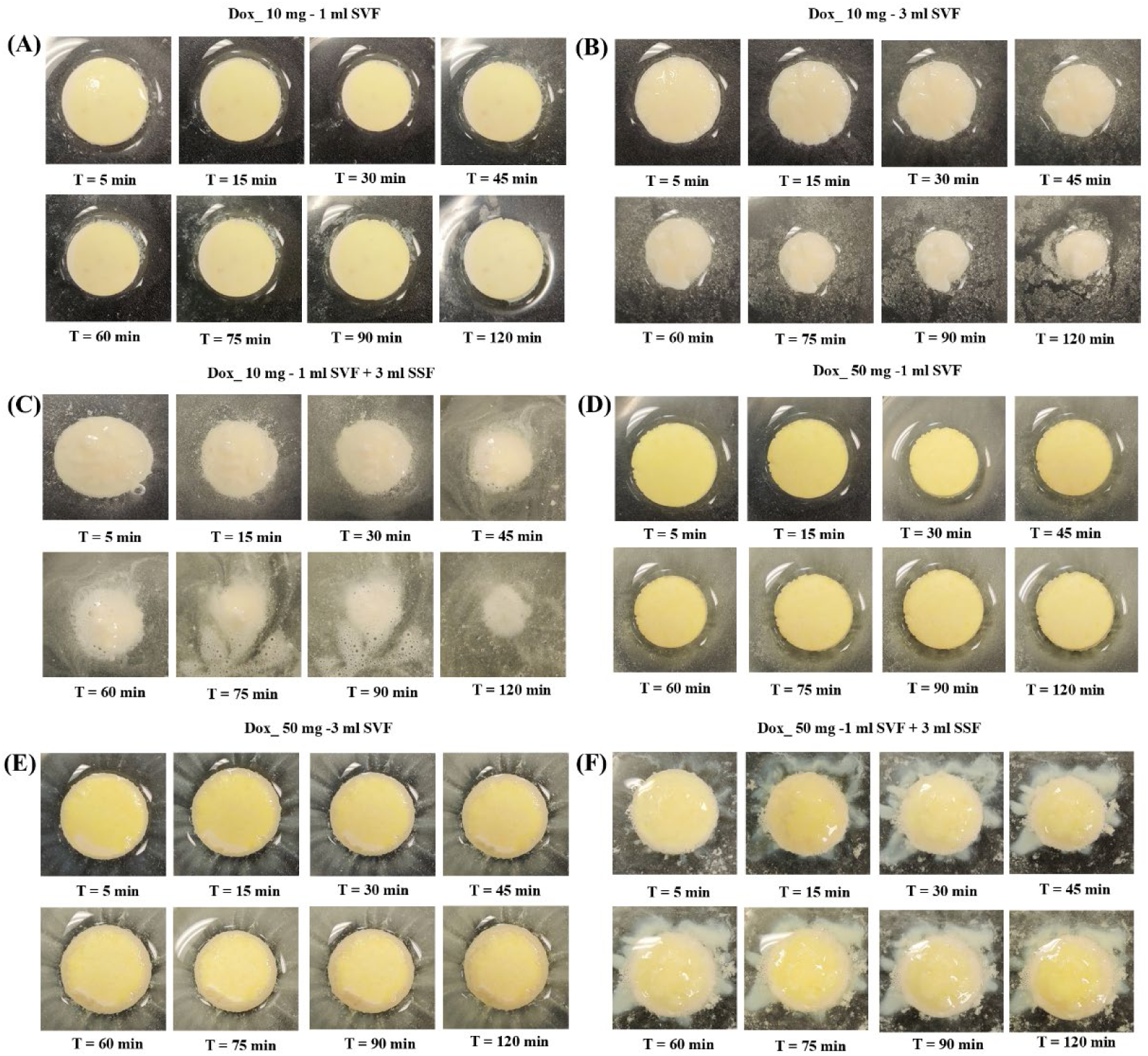
Visual assessments of inserts in vitro disintegration at 37°C: (A) 10 mg DOX insert in 1 mL SVF at pH ∼4.3; (B) 10 mg DOX insert in 3 mL SVF at pH ∼4.3; (C) 10 mg DOX insert in 4 mL simulant mixture at pH ∼7.2; (D) 50 mg DOX insert in 1 mL SVF at pH ∼4.3; (E) 50 mg DOX insert in 3 mL SVF at pH ∼4.3; (F) 50 mg DOX insert in 4 mL simulant mixture at pH ∼7.2.

**Table 2.** Physicochemical characteristics of DOX inserts.

| Parameters | D1<br>(10 mg DOX)* | D2<br>(50 mg<br>DOX)* | D3<br>(100 mg<br>DOX)* |
| --- | --- | --- | --- |
| Physical appearance (n = 10) | Off white to off yellow, round | Off yellow, round | Off yellow, round |
| Hardness (kg) ± SD (n = 10) | 7.3 ± 0.6 | 7.5 ± 0.4 | 6.7 ± 0.5 |
| Friability (average % weight loss) (n = 13) | 0.502 | 0.511 | 0.357 |
| Drug assay (%) ± SD (n = 3) | 99.13 ± 1.73 | 100.25 ± 1.06 | 99.90 ± 1.25 |
| Moisture content (%) ± SD (n = 5) | 0.638 ± 0.004 | 0.719 ± 0.005 | 0.983 ± 0.070 |
| Osmolality (mOsm/kg) in 5 mL of SVF (pH ~4.3) (n = 3) | 535 ± 7 | 519 ± 11 | 513 ± 9 |
| Disintegration time (min) ± SD (n = 6) in acetate buffer at pH ~4.5 | 9.15 ± 0.71 | 8.10 ± 0.44 | 9.13 ± 0.43 |
| Cumulative drug release (%) ± SD at 30 min of insert dissolution (n = 6) in acetate buffer at pH ~4.5 | 100.0 ± 0.9 | 87.7 ± 3.9 | 97.5 ± 1.0 |
| Cumulative drug release (%) ± SD at 60 min of insert dissolution (n = 6) in acetate buffer at pH ~4.5 | 102.5 ± 1.5 | 99.6 ± 2.4 | 104.1 ± 2.6 |

\* Inserts containing DH equivalent to 10 (D1), 50 (D2), and 100 (D3) mg of DOX.

### Stability Testing of DOX Inserts

The DOX inserts maintained their physical appearance, drug assay, hardness, friability, and disintegration under controlled room-temperature (25°C/60% RH) and accelerated (40°C/75% RH) stability conditions, as tested for 6 months **(Table 3)**. The osmolality and moisture content of the inserts were also maintained below <550 mOsm/kg and <2%, respectively (tested at initial and 6-month time points). Under the USP dissolution testing conditions, inserts demonstrated comparable DOX release (more than 85% at 30 min of analysis) at both initial (time zero) and 6-month time points under 25°C/60% RH and 40°C/75% RH **(Fig. 3)**.

**Fig. 3.**
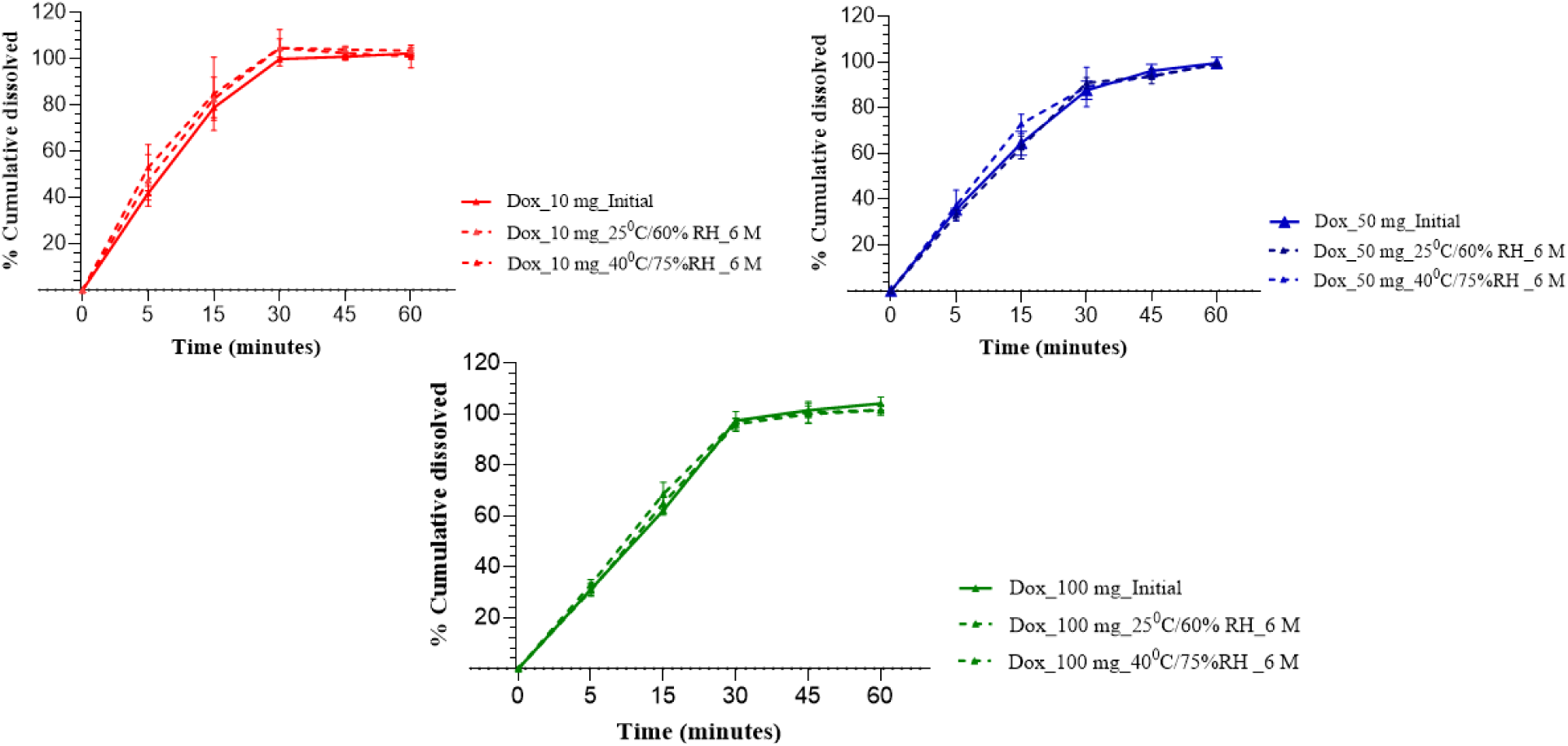
Cumulative (%) DOX in vitro release from inserts (10, 50, and 100 mg DOX) in acetate buffer at pH ∼4.5, for inserts stored at 25°C/60% RH and 40°C/75% RH for 6 months. The results are reported as the mean ± SD of six inserts tested at each time point.

**Table 3:**
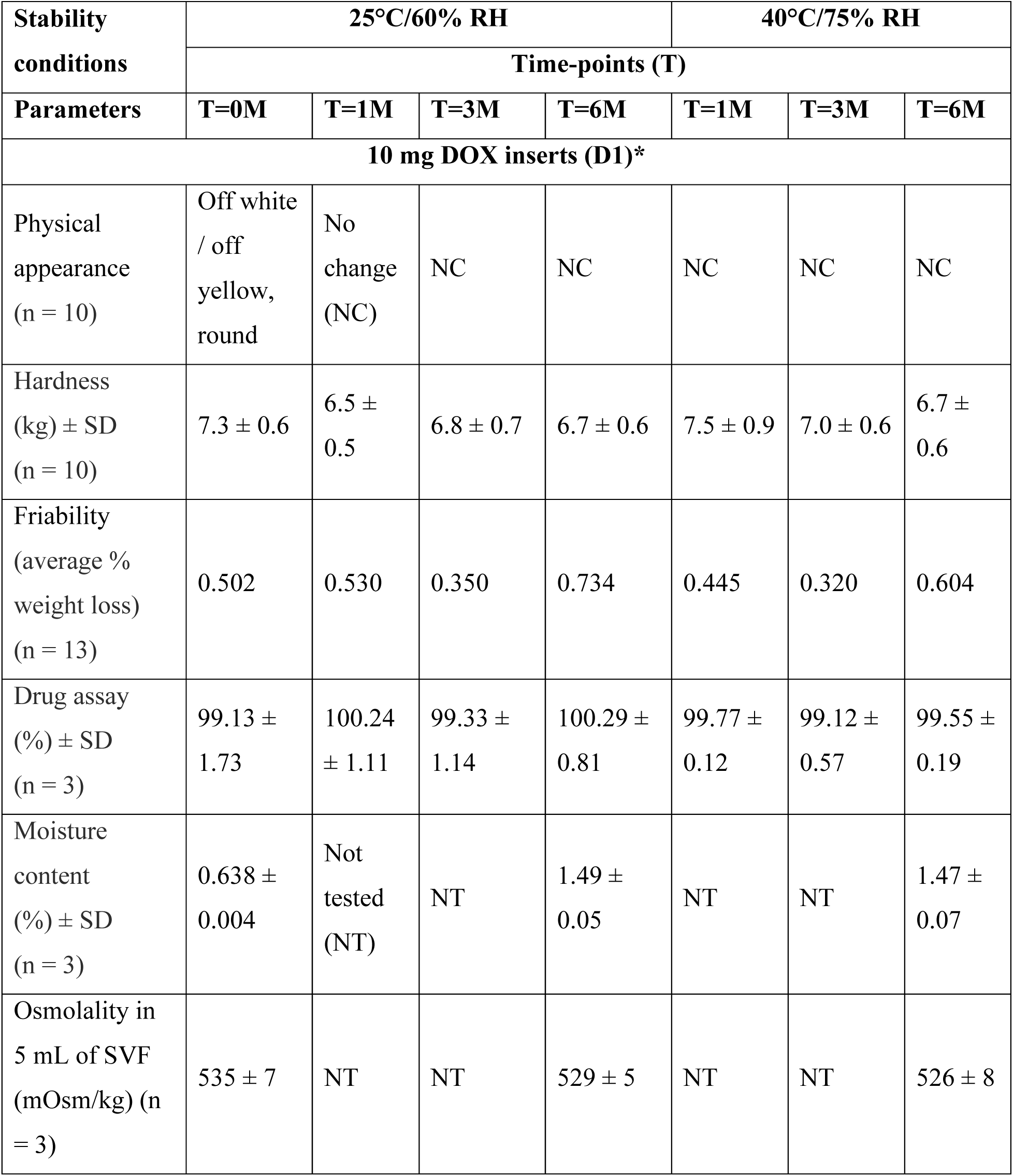
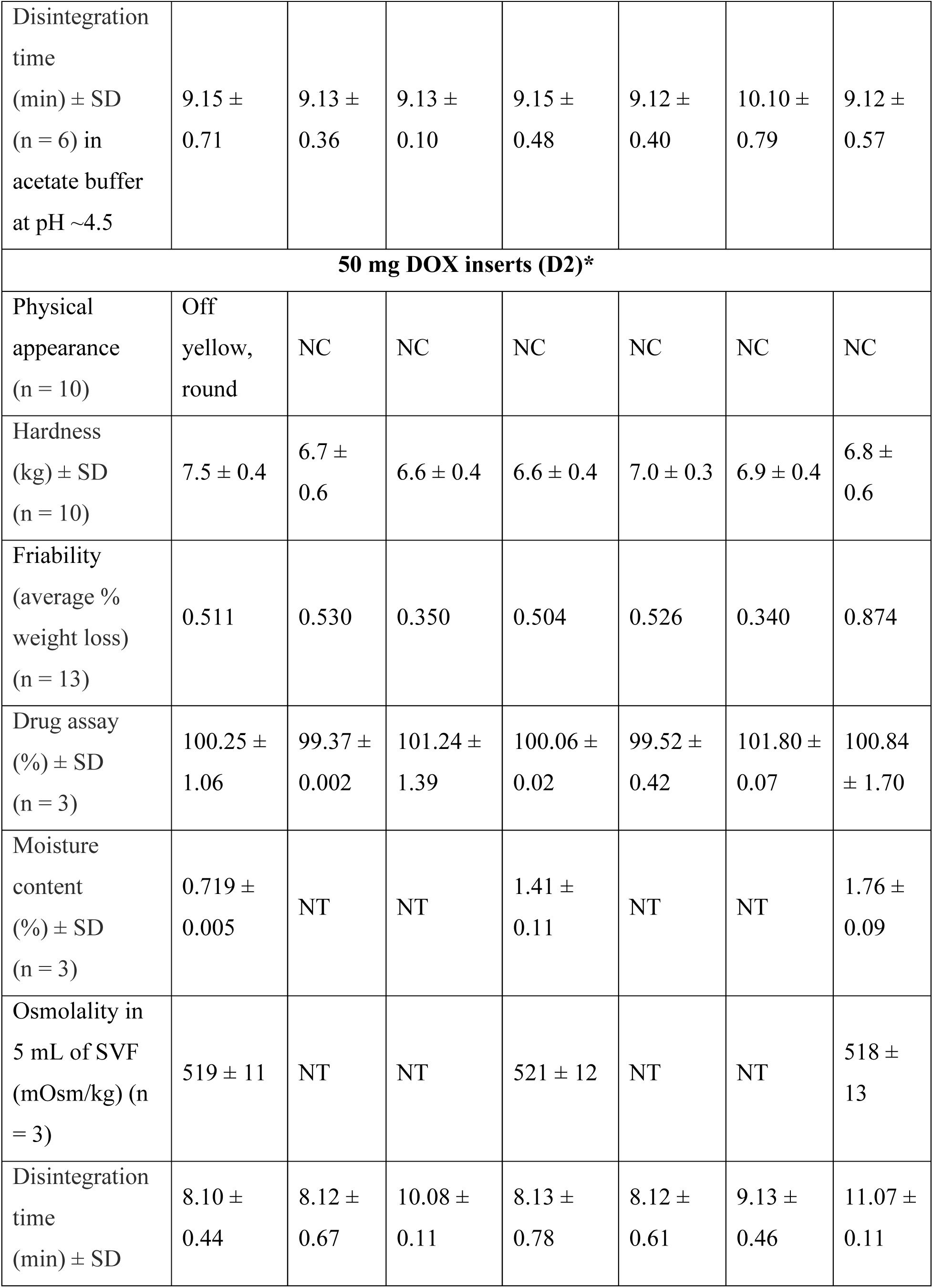
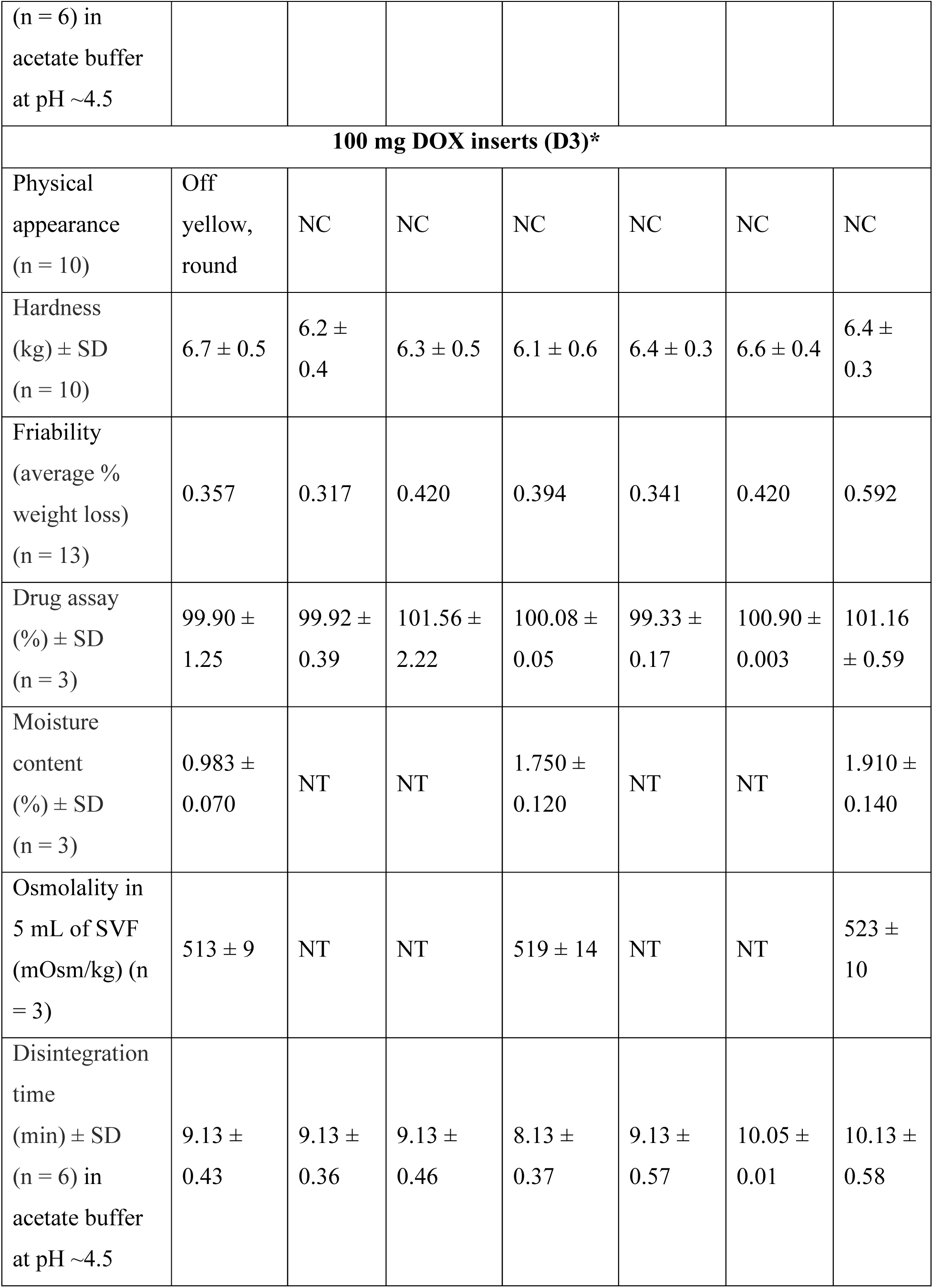
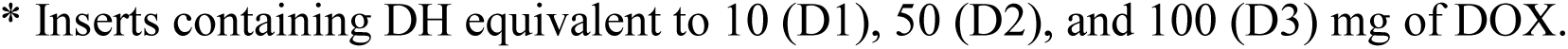
Stability testing of DOX inserts.

### TED Insert Formulation Development and Characterization

The representative HPLC chromatogram of TAF, EVG, and DOX (DH) with RT of ∼5.35, 8.47, and 5.13 min, respectively, is shown in **Supplementary Fig. 2.** The calibration curves for TAF, EVG, and DOX (DH) in the concentration range of 3.1-225 µg/mL were linear with R^2^ of >0.999. The composition of TED inserts **(Fig. 4)** is provided in **Table 4**.

**Fig. 4.**
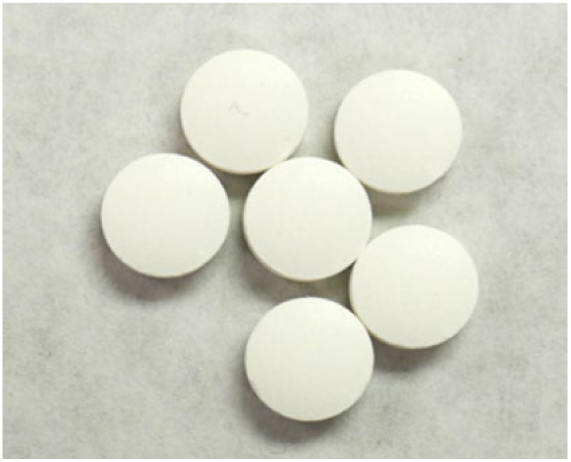
TED inserts (500 mg) containing TAF free base (20 mg), EVG (16 mg), and DOX (10 mg).

**Table 4:** Composition of TED inserts.

| Materials | % w/w | Weight (mg) |
| --- | --- | --- |
| TAF | 4.48 | 22.40* |
| EVG | 3.20 | 16.00 |
| DOX hyclate (DH) | 2.31 | 11.55* |
| Lactose anhydrous | 53.71 | 268.55 |
| Povidone (K-29/32) | 4.00 | 20.00 |
| Poloxamer 188 | 2.00 | 10.00 |
| PEG 8000 | 15.00 | 75.00 |
| Mannitol | 13.80 | 69.00 |
| Magnesium stearate | 1.500 | 7.50 |
| Total | 100 | 500 |

\* Equivalent to 20 mg of TAF free base and 10 mg of DOX.

The physicochemical characteristics of TED inserts are summarized in **Table 5**. The inserts showed drug assays within 95-105% of the initial drug content, hardness within 5-8 kg, friability <1%, osmolality <550 mOsm/kg, and moisture content <2%. Under the USP testing method, complete disintegration of the inserts was observed within ∼10 min, achieving over 95% cumulative release of TAF, EVG, and DOX within 60 min of dissolution.

**Table 5.** Physicochemical characteristics of TED inserts.

| Parameters | Values |
| --- | --- |
| Physical appearance (n = 10) | Off white to off yellow, round |
| Hardness (kg) ± SD (n = 10) | 5.60 ± 0.30 |
| Friability (average % weight loss) (n = 13) | 0.59 |
| Drug assay (%) ± SD (n = 3) | TAF: 99.70 ± 0.30; EVG: 100.50 ± 0.30; DOX (DH): 100.50 ± 1.30 |
| Moisture content (%) ± SD (n = 5) | 1.04 ± 0.20 |
| Disintegration time (min) ± SD (n = 6) in acetate buffer at pH ~4.5 | 8.70 ± 0.61 |
| Cumulative drug release (%) ± SD at 60 min of insert dissolution (n = 3) in acetate buffer at pH ∼4.5 | TAF: $97.42 \pm 1.48$; EVG: $95.76 \pm 0.51$; DOX (DH): $95.26 \pm 1.08$ |
| Osmolality (mOsm/kg) in 5 mL of SVF (pH ∼4.3) (n = 3) | $513 \pm 10$ |

The 1-month pilot stability data for TED inserts at 40°C/75% RH, compared to the initial (time zero), showed that inserts maintained their physical appearance (off-white to off-yellow, round), hardness (6.62 ± 0.61 kg, n = 10), and friability (1.12, n = 13). A friability value greater than 1 indicates a potential structural weakness of the TED inserts under high temperatures and humidity, suggesting the need to optimize the binder type and concentration. There were no significant changes observed in the drug assays (TAF: 98.63 ± 0.9; EVG: 99.13 ± 1.73; DOX: 99.75 ± 0.21, n = 3) nor dissolution profiles **(Fig. 5)** compared to initial (t = 0) data.

**Fig. 5.**
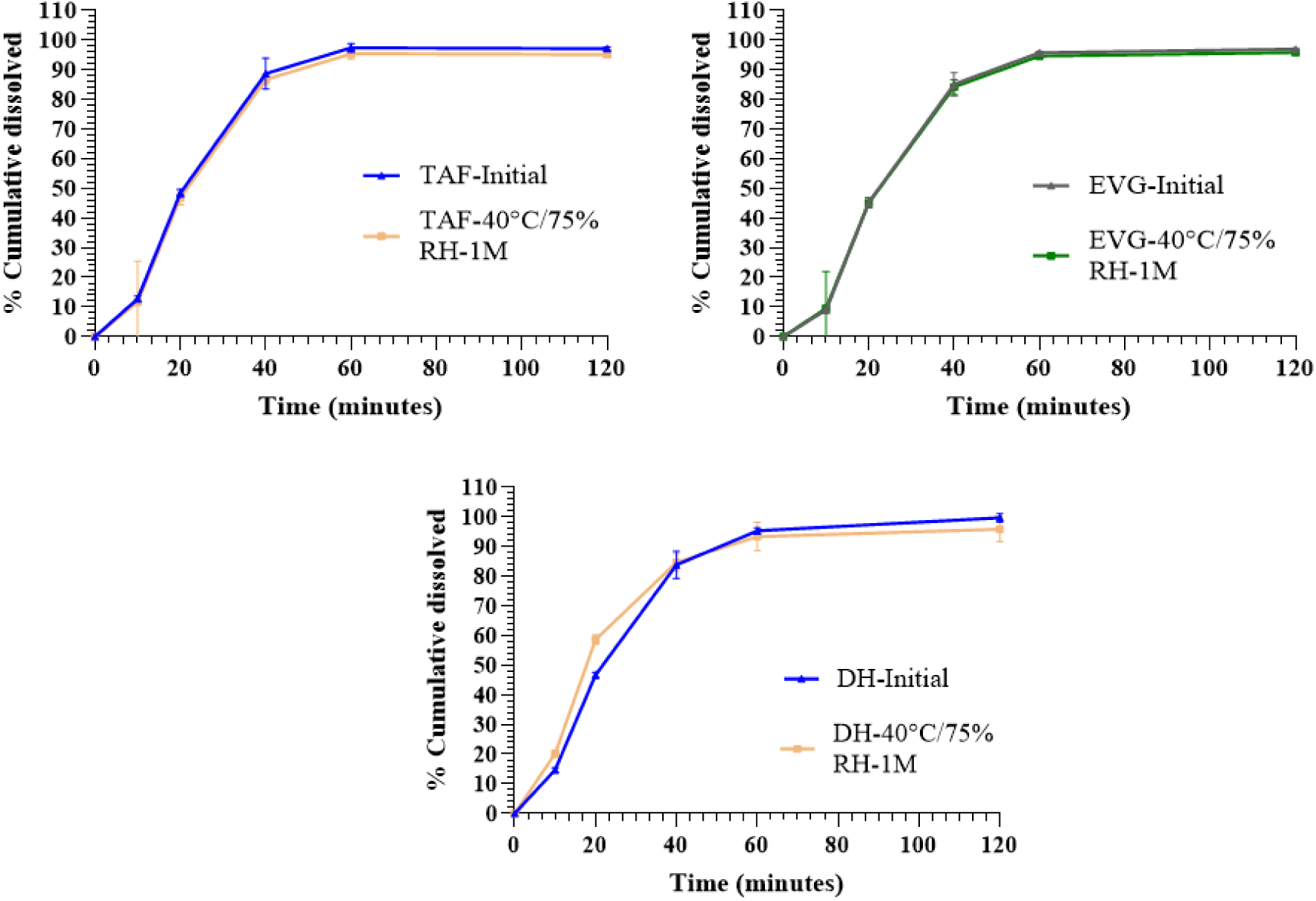
Cumulative (%) TAF, EVG, and DOX in vitro release from TED inserts in acetate buffer at pH ∼4.5, for inserts stored at 40°C/75% RH for 1 month, compared to initial (t = 0) data. The results are reported as the mean ± SD of six inserts tested at each time point.

## DISCUSSION

We have developed DOX-only and TED inserts for the prevention of STIs (bacterial and bacterial/viral). Building on experience from our previously developed TAF/EVG inserts already tested in Phase I clinical trials, the direct compression process was chosen for DOX and TED inserts manufacturing due to its simplicity and ease of scale-up, thereby making it more efficient and cost-effective [19]. Additionally, this method yields inserts with superior mechanical strength, which is essential during handling, transportation, and packaging. This increased durability helps improve product stability and extends shelf life. Regarding the rationale for selecting specific insert excipients, lactose and mannitol were used as diluents to provide the required volume and weight for the inserts. Selected for their excellent flowability, low hygroscopicity, and high compressibility, they support consistent die filling, uniform weight, and mechanical integrity during compression [34–36]. Lactose also contributes to water solubility, whereas mannitol facilitates the formation of inserts that disintegrate rapidly, thereby ensuring rapid drug release. Povidone (polyvinylpyrrolidone, PVP) was included as a binder to improve mechanical strength and cohesiveness by forming hydrogen bonds with excipients and drugs, preventing premature disintegration during manufacturing and storage [37,38]. PEG 8000, a water-soluble polymer, enhances dissolution and disintegration while also acting as a binder, improving interparticle cohesion and mechanical stability [39]. This dual role ensures that the inserts remain mechanically stable during handling and packaging while dissolving quickly upon contact with physiological fluids. Poloxamer 188, an amphiphilic nonionic copolymer, functions as a disintegration and dissolution enhancer by lowering interfacial tension and facilitating water uptake [40]. It also provides lubrication, reducing die-wall friction and improving processability. Magnesium stearate, an efficient lubricant at low concentrations, coats particle surfaces to minimize friction and stick to die walls and punches, enhancing flowability and ensuring uniform die filling and compressibility [41,42].

The physicochemical properties of topical inserts are crucial in determining their performance, as they affect disintegration, dissolution, drug release, safety upon mucosal application, and overall formulation stability. The drug content assay of DOX in DOX-only and TAF/EVG/DOX in TED inserts was maintained within 95-105% of the labeled content, demonstrating a robust manufacturing process that ensures batch-to-batch reproducibility for accurate dosing. Both DOX and TED inserts were designed with a target hardness of 5-8 kg and friability of <1% to ensure structural integrity, which were achieved. Results from our past insert projects suggest that this range offers a balance between sufficient strength to prevent chipping or breaking and adequate porosity and friability to allow for faster disintegration and drug release upon use. Although tested for only one month under accelerated conditions, a friability value of >1 (1.12) for the TED inserts highlights the need to optimize the binder type and concentration in the formulation to enhance stability during long-term storage. Nevertheless, additional long-term stability studies under both room temperature and accelerated conditions are required to validate these observations.

The disintegration of solid dosage forms, such as inserts, typically begins with the absorption of water into the core, leading to pore formation that accelerates dissolution and facilitates drug release [43]. This process can occur through bulk or surface erosion, or a combination of both, and is influenced by several factors, including the properties of the drug, excipients, and dissolution medium [44]. Under USP testing conditions, the in vitro disintegration of both DOX and TED inserts started within 5 min (completed in ∼10 min) and released >95% of DOX (in all three strengths of DOX-only inserts) or TAF/EVG/DOX (in TED insert) within 60 min of initiation of insert dissolution. Visual assessments of the DOX-only inserts (10 and 50 mg strengths) revealed that dissolution behavior was influenced by the volume and composition of the dissolution medium and the amount of drug (used in acidic hyclate salt form), which, together, may have influenced DOX solubility. Observations revealed that the 10 mg DOX insert, when tested in 3 mL of SVF, primarily underwent surface erosion, a favorable mechanism for achieving more consistent and predictable drug release, as it relies on continuous surface-fluid interaction and is independent of the internal matrix volume [45]. In contrast, minimal surface erosion was seen for the 10 mg insert in a 1 mL volume. Similarly, the 50 mg DOX inserts showed minimal surface erosion under both volumes, suggesting that higher drug loading and/or limited fluid volume may prevent enough surface interaction and reduce effective erosion. A mixture of vaginal and seminal fluid simulants (1:4% v/v) with a higher pH (∼7.2) seemed to improve disintegration. It is worth noting that both 10 mg and 50 mg DOX inserts, when tested vaginally in monkeys, released DOX, achieving high and long-lasting vaginal concentrations that prevented chlamydial infection [30]. In a previous CONRAD Phase I clinical study of tenofovir and emtricitabine vaginal inserts, although insert dissolution proceeded at a slower-than-expected rate, drug release commenced as soon as the inserts were imbibed in fluid (manuscript in preparation). Furthermore, in previous studies in NHPs and women, TAF/EVG inserts with a very similar composition to the ones reported in this paper rapidly disintegrated in the vagina, achieving high concentrations of TAF and EVG at the first timepoint of sample collection, 2 and 4 h, respectively, after insertion [21,46]. Future research should explore the dissolution behaviors of DOX and TED inserts under physiologically relevant and clinical conditions.

Moisture content is a key factor impacting the long-term stability, mechanical strength, and disintegration behavior of solid dosage form inserts. For DOX inserts, silica gel desiccant in the HDPE packaging bottles effectively maintained the inserts’ moisture levels, including under ambient and accelerated stability conditions, as tested for 6 months. The moisture content, although still below 2%, increased compared to the time-zero data. However, this rise did not affect the disintegration or dissolution profiles of the DOX inserts. These findings suggest that using a higher amount of silica gel desiccant (e.g., 3 g), as also employed for the clinically advanced TAF/EVG insert [19], may help maintain lower moisture levels in the DOX and TED inserts over an extended period. Osmolality is also a key physicochemical parameter in the design of vaginal formulations, as it influences mucosal tolerability and acceptability [47]. In 5 mL of SVF, both DOX and TED inserts exhibited osmolalities under 550 mOsm/kg, well below the WHO provisional limit of 1200 mOsm/kg for intravaginal products [48]. The inserts were also designed to have a longer shelf life for storage. Based on 6-month stability data for DOX-only inserts under accelerated and long-term storage conditions, the physicochemical attributes of the inserts were maintained relative to time-zero data and are projected to confer a shelf life of over 2 years if stored at or below 25°C/60% RH. The TED inserts were tested for only 1-month at 40°C/75% RH. Although the stability results appear promising, further long-term testing is necessary to confirm their shelf life and optimal storage conditions.

## CONCLUSION AND FUTURE DIRECTIONS

We described the development of two distinct topical inserts: one containing DOX only and another combining DOX with TAF and EVG (TED insert). The DOX insert is designed to prevent bacterial STIs such as chlamydia, gonorrhea, and syphilis, while the TED insert offers a multi-purpose prevention approach targeting both bacterial STIs and HIV-1. These topical inserts aim to broaden prevention options by delivering high drug concentrations locally (vaginally and/or rectally), directly at the primary infection site. This targeted, on-demand application helps reduce systemic exposure and side effects while reducing the number of daily doses required, addressing unmet needs in current prevention strategies. Further investigations will be necessary to assess the potential for antimicrobial resistance with intermittent topical administration and how it compares to that seen with standard-of-care oral DOX [49,50]. Furthermore, as these topical inserts can be used vaginally or rectally, before or after coitus, they provide a highly convenient, portable, discreet prevention option for both women and men that is unlike any other HIV or STI PrEP/PEP methods currently available. The inserts have several promising features. In addition, they use FDA-approved, Generally Recognized As Safe (GRAS) excipients and are produced through a simple, cost-effective manufacturing process that is easily scalable and transferable for regional manufacturing globally. The final products demonstrate strong mechanical integrity, appropriate disintegration and drug release profiles, and long-term stability.

We have demonstrated that the topical insert platform used for the clinically advanced TAF/EVG insert provides similarly compatible and stable formulations for the DOX and TED inserts, demonstrating promising characteristics; however, additional preclinical studies in animal models, particularly focused on efficacy, are necessary. Testing of the DOX insert in NHPs is currently underway in collaboration with the CDC to assess PK and efficacy [30]. Preliminary findings have shown that NHPs that received DOX inserts [50 mg or 10 mg] were protected against vaginal CT infection, whereas almost all controls (5/6) became infected. Vaginal administration of a single 10 mg DOX insert resulted in DOX concentrations in vaginal swab eluates exceeding the MIC_90_ of DOX against CT (0.064 μg/mL) for ≥10 days, with undetectable systemic DOX exposure. Furthermore, DOX levels were higher and lasted longer than vaginal concentrations achieved after a single oral dose (200 mg) of DOX [30], currently recommended for prevention (PEP) of chlamydial infection [26,51]. Higher and longer-lasting concentrations of DOX in the vaginal compartment may be critical to demonstrating efficacy in women [15]. The TED inserts require slight optimization to ensure their long-term stability before advancing to similar safety, PK, and efficacy evaluations in NHPs. Having completed scale-up, clinical manufacturing, and testing of a similar insert (TAF/EVG: 20 mg/16 mg) under current Good Manufacturing Practice (cGMP) conditions, we anticipate that the manufacturing process for the DOX and TED inserts can be readily scaled and transferred. Moreover, the topical insert platform is adaptable and can deliver a range of antiviral and antibacterial agents, either vaginally or rectally, to further support the development of multipurpose prevention products for single-dose protection against STIs. Plans for testing the TED insert in a first-in-human Phase I clinical trial are underway.

## ACKNOWLEDGMENTS

The authors thank Gilead Sciences, Inc. (Foster City, CA) for providing EVG drug substance for the TED insert development activities.

## CRediT AUTHORSHIP CONTRIBUTION STATEMENT

**Vivek Agrahari:** Conceptualization, Methodology, Validation, Formal analysis, Investigation, Writing – Original Draft, Writing – Review and Editing, Visualization. **M. Melissa Peet:** Conceptualization, Methodology, Validation, Formal analysis, Investigation, Writing – Original Draft, Writing – Review and Editing, Visualization. **Jasmin Monpara:** Methodology, Validation, Formal analysis, Resources, Data curation, Writing – Review and Editing. **Rijo John:** Methodology, Validation, Formal analysis, Resources, Data curation, Writing – Review and Editing. **Pardeep K. Gupta:** Supervision, Project administration, Data curation, Writing – Review and Editing. **Sriramakamal Jonnalagadda:** Supervision, Project administration, Data curation, Writing – Review and Editing. **Meredith R. Clark:** Conceptualization, Visualization, Writing – Review and Editing. **Gustavo F. Doncel:** Conceptualization, Visualization, Supervision, Funding acquisition, Writing – Review and Editing.

## DATA AVAILABILITY

The datasets generated during and/or analyzed during the current study are available from the corresponding author on reasonable request.

## REFERENCES

1. Fact Sheet: Global HIV & AIDS statistics [Internet]. UNAIDS; 2025 Jul 10 [cited 2025 Sep 22]. Available from: https://www.unaids.org/en/resources/fact-sheet.

2. Fact Sheet: Sexually transmitted infections (STIs) [Internet]. WHO; 2025 Sep 10 [cited 2025 Oct 7]. Available from: https://www.who.int/news-room/fact-sheets/detail/sexually-transmitted-infections-(stis).

3. Ward H, Rönn M. Contribution of sexually transmitted infections to the sexual transmission of HIV. Curr Opin HIV AIDS. 2010;5(4):305–10.

4. National Academies of Sciences E, Medicine, Health, Medicine D, Board on Population H, Public Health P, et al. In: Crowley JS, Geller AB, Vermund SH, editors. Sexually Transmitted Infections: Adopting a Sexual Health Paradigm. Washington (DC): National Academies Press (US) Copyright 2021 by the National Academy of Sciences. All rights reserved.; 2021.

5. Cohen MS, Council OD, Chen JS. Sexually transmitted infections and HIV in the era of antiretroviral treatment and prevention: the biologic basis for epidemiologic synergy. J Int AIDS Soc. 2019;22 Suppl 6(Suppl Suppl 6):e25355.

6. Mwatelah R, McKinnon LR, Baxter C, Abdool Karim Q, Abdool Karim SS. Mechanisms of sexually transmitted infection-induced inflammation in women: implications for HIV risk. J Int AIDS Soc. 2019;22 Suppl 6(Suppl Suppl 6):e25346.

7. Mayer KH, Venkatesh KK. Interactions of HIV, other sexually transmitted diseases, and genital tract inflammation facilitating local pathogen transmission and acquisition. Am J Reprod Immunol. 2011;65(3):308–16.

8. Preventing Sexual Transmission of HIV. HIV.gov. Updated September 8, 2025. Accessed October 7, 2025. https://www.hiv.gov/hiv-basics/hiv-prevention/reducing-sexual-risk/preventing-sexual-transmission-of-hiv.

9. Yap PK, Loo Xin GL, Tan YY, Chellian J, Gupta G, Liew YK, et al. Antiretroviral agents in pre-exposure prophylaxis: emerging and advanced trends in HIV prevention. J Pharm Pharmacol. 2019;71(9):1339–52.

10. Riddell Jt, Amico KR, Mayer KH. HIV Preexposure Prophylaxis: A Review. Jama. 2018;319(12):1261–8.

11. Chou R, Spencer H, Bougatsos C, Blazina I, Ahmed A, Selph S. Preexposure Prophylaxis for the Prevention of HIV: Updated Evidence Report and Systematic Review for the US Preventive Services Task Force. Jama. 2023;330(8):746–63.

12. Boschiero MN, Sansone NMS, Matos LR, Marson FAL. Efficacy of Doxycycline as Preexposure and/or Postexposure Prophylaxis to Prevent Sexually Transmitted Diseases: A Systematic Review and Meta-Analysis. Sexually Transmitted Diseases. 2025;52(2).

13. Molina JM, Charreau I, Chidiac C, Pialoux G, Cua E, Delaugerre C, et al. Post-exposure prophylaxis with doxycycline to prevent sexually transmitted infections in men who have sex with men: an open-label randomised substudy of the ANRS IPERGAY trial. Lancet Infect Dis. 2018;18(3):308–17.

14. Bolan RK, Beymer MR, Weiss RE, Flynn RP, Leibowitz AA, Klausner JD. Doxycycline prophylaxis to reduce incident syphilis among HIV-infected men who have sex with men who continue to engage in high-risk sex: a randomized, controlled pilot study. Sex Transm Dis. 2015;42(2):98–103.

15. Stewart J, Oware K, Donnell D, Violette LR, Odoyo J, Soge OO, et al. Doxycycline Prophylaxis to Prevent Sexually Transmitted Infections in Women. N Engl J Med. 2023;389(25):2331–40.

16. Tanne JH. US public health agency recommends prophylactic doxycycline to reduce STIs. Bmj. 2023;383:2339.

17. Dietrich JJ, Ahmed N, Webb EL, Tshabalala G, Hornschuh S, Mulaudzi M, et al. A multi-country cross-sectional study to assess predictors of daily versus on-demand oral pre-exposure prophylaxis in youth from South Africa, Uganda and Zimbabwe. J Int AIDS Soc. 2022;25(8):e25975.

18. Friedland BA, Plagianos M, Savel C, Kallianes V, Martinez C, Begg L, et al. Women Want Choices: Opinions from the Share.Learn.Shape Global Internet Survey About Multipurpose Prevention Technology (MPT) Products in Development. AIDS Behav. 2023;27(7):2190–204.

19. Agrahari V, Peet MM, Chandra N, Ramalingam P, Gupta PK, Jonnalagadda S, et al. Formulation development of dual-compartment topical inserts combining tenofovir alafenamide and elvitegravir for flexible on-demand HIV prevention. J Control Release. 2025;377:842–54.

20. Peet MM, Agrahari V, Clark MR, Doncel GF. Preclinical and Early Clinical Development of Tenofovir Alafenamide/Elvitegravir Topical Inserts for Effective On-Demand Vaginal and Rectal HIV Prevention. Pharmaceutics. 2024;16(3).

21. Thurman AR, Ouattara LA, Yousefieh N, Anderson PL, Bushman LR, Fang X, et al. A phase I study to assess safety, pharmacokinetics, and pharmacodynamics of a vaginal insert containing tenofovir alafenamide and elvitegravir. Front Cell Infect Microbiol. 2023;13:1130101.

22. Riddler SA, Kelly CW, Hoesley CJ, Ho KS, Piper JM, Edick S, et al. A Phase 1 Clinical Trial to Assess the Safety and Pharmacokinetics of a Tenofovir Alafenamide/Elvitegravir Insert Administered Rectally for HIV Prevention. J Infect Dis. 2024;230(3):696–705.

23. Peet MM, Agrahari V, Anderson SM, Hanif H, Singh ON, Thurman AR, et al. Topical Inserts: A Versatile Delivery Form for HIV Prevention. Pharmaceutics. 2019;11(8).

24. Agwuh KN, MacGowan A. Pharmacokinetics and pharmacodynamics of the tetracyclines including glycylcyclines. Journal of Antimicrobial Chemotherapy. 2006;58(2):256–65.

25. Saivin S, Houin G. Clinical Pharmacokinetics of Doxycycline and Minocycline. Clinical Pharmacokinetics. 1988;15(6):355–66.

26. Haaland RE, Fountain J, Edwards TE, Dinh C, Martin A, Omoyege D, et al. Pharmacokinetics of single dose doxycycline in the rectum, vagina, and urethra: implications for prevention of bacterial sexually transmitted infections. eBioMedicine. 2024;101.

27. Ljubin-Sternak S, Meštrović T. Antimicrobial Susceptibility Testing in Chlamydia trachomatis: The Current State of Evidence and a Call for More National Surveillance Studies. Applied Sciences. 2025;15(8):4322.

28. Zheng XL, Xu WQ, Liu JW, Zhu XY, Chen SC, Han Y, et al. Evaluation of Drugs with Therapeutic Potential for Susceptibility of Neisseria Gonorrhoeae Isolates from 8 Provinces in China from 2018. Infect Drug Resist. 2020;13:4475–86.

29. Edmondson DG, Wormser GP, Norris SJ. In Vitro Susceptibility of Treponema pallidum subsp. pallidum to Doxycycline. Antimicrob Agents Chemother. 2020;64(10).

30. Garber DA, Peet MM, Jia H, Priode JH, Mitchell J, Dinh C, Edwards E, Haaland R, Agrahari V, Clark MR, Vishwanathan SA, McNicholl J, Doncel GF, Heneine W. Topical Doxycycline Inserts Show High Efficacy Against Vaginal Chlamydia Acquisition in Macaques [abstract]. In: Proceedings of (T-05) DoxyPEP Reflections [Internet]. CROI: 2025 Mar 9-12; San Francisco, CA. Available from: https://www.croiconference.org/abstract/577-2025/.

31. Owen DH, Katz DF. A vaginal fluid simulant. Contraception. 1999;59(2):91–5.

32. Owen DH, Katz DF. A review of the physical and chemical properties of human semen and the formulation of a semen simulant. J Androl. 2005;26(4):459–69.

33. Stability Testing of New Drug Substances and Products Q1A(R2). International Council for Harmonisation of Technical Requirements for Pharmaceuticals for Human Use. November 2003.

34. Bolhuis GK, Anthony Armstrong N. Excipients for Direct Compaction—an Update. Pharmaceutical Development and Technology. 2006;11(1):111–24.

35. Jivraj M, Martini LG, Thomson CM. An overview of the different excipients useful for the direct compression of tablets. Pharmaceutical Science & Technology Today. 2000;3(2):58–63.

36. Ohrem HL, Schornick E, Kalivoda A, Ognibene R. Why is mannitol becoming more and more popular as a pharmaceutical excipient in solid dosage forms? Pharmaceutical Development and Technology. 2014;19(3):257–62.

37. Kurakula M, Rao GSNK. Pharmaceutical assessment of polyvinylpyrrolidone (PVP): As excipient from conventional to controlled delivery systems with a spotlight on COVID-19 inhibition. Journal of Drug Delivery Science and Technology. 2020;60:102046.

38. Kouass S, Bejaoui M, Galai H, Touati F. Multifunctional Roles of PVP as a Versatile Biomaterial in Solid State. In: Ahmad U, editor. Dosage Forms - Innovation and Future Perspectives. London: IntechOpen; 2021.

39. Shah RC, Raman PV, Sheth PV. Polyethylene Glycol as a Binder for Tablets. Journal of Pharmaceutical Sciences. 1977;66(11):1551–2.

40. Kibbe AH. Handbook of pharmaceutical excipients. 3rd ed: American Pharmaceutical Association,Pharmaceutical Press; 2000.

41. Morin G, Briens L. The Effect of Lubricants on Powder Flowability for Pharmaceutical Application. AAPS PharmSciTech. 2013;14(3):1158–68.

42. Wang J, Wen H, Desai D. Lubrication in tablet formulations. Eur J Pharm Biopharm. 2010;75(1):1–15.

43. Desai PM, Liew CV, Heng PWS. Review of Disintegrants and the Disintegration Phenomena. Journal of Pharmaceutical Sciences. 2016;105(9):2545–55.

44. Markl D, Zeitler JA. A Review of Disintegration Mechanisms and Measurement Techniques. Pharmaceutical Research. 2017;34(5):890–917.

45. Tamada JA, Langer R. Erosion kinetics of hydrolytically degradable polymers. Proc Natl Acad Sci U S A. 1993;90(2):552–6.

46. Dobard CW, Peet MM, Nishiura K, Holder A, Dinh C, Mitchell J, et al. Single dose topical inserts containing tenofovir alafenamide fumarate and elvitegravir provide pre- and post-exposure protection against vaginal SHIV infection in macaques. EBioMedicine. 2022;86:104361.

47. Ayehunie S, Wang YY, Landry T, Bogojevic S, Cone RA. Hyperosmolal vaginal lubricants markedly reduce epithelial barrier properties in a three-dimensional vaginal epithelium model. Toxicol Rep. 2018;5:134–40.

48. Use and procurement of additional lubricants for male and female condoms: WHO/UNFPA/FHI360. World Health Organization. Advisory note. November 1, 2012. Accessed October 7, 2025. https://iris.who.int/server/api/core/bitstreams/c0d0726b-e941-4d63-9647-fe2c2a52438a/content.

49. Truong R, Tang V, Grennan T, Tan DHS. A systematic review of the impacts of oral tetracycline class antibiotics on antimicrobial resistance in normal human flora. JAC-Antimicrobial Resistance. 2022;4(1).

50. Soge OO, Thibault CS, Cannon CA, McLaughlin SE, Menza TW, Dombrowski JC, et al. Potential Impact of Doxycycline Post-Exposure Prophylaxis on Tetracycline Resistance in Neisseria gonorrhoeae and Colonization With Tetracycline-Resistant Staphylococcus aureus and Group A Streptococcus. Clin Infect Dis. 2025;80(6):1188–96.

51. Bachmann LH, Barbee LA, Chan P, Reno H, Workowski KA, Hoover K, Mermin J, Mena L. CDC Clinical Guidelines on the Use of Doxycycline Postexposure Prophylaxis for Bacterial Sexually Transmitted Infection Prevention, United States, 2024. U.S. MMWR Recomm Rep 2024;73(2).

