## Supplementary file for "Formulation Development of Topical Inserts Containing Doxycycline and Doxycycline Combined with Tenofovir Alafenamide and Elvitegravir for the Prevention of Sexually Transmitted Infections"

### SUPPLEMENTARY FIGURES

**Supplementary Figure 1:** The representative HPLC chromatogram of DOX (DH) with a retention time of ~15.79 min**.**

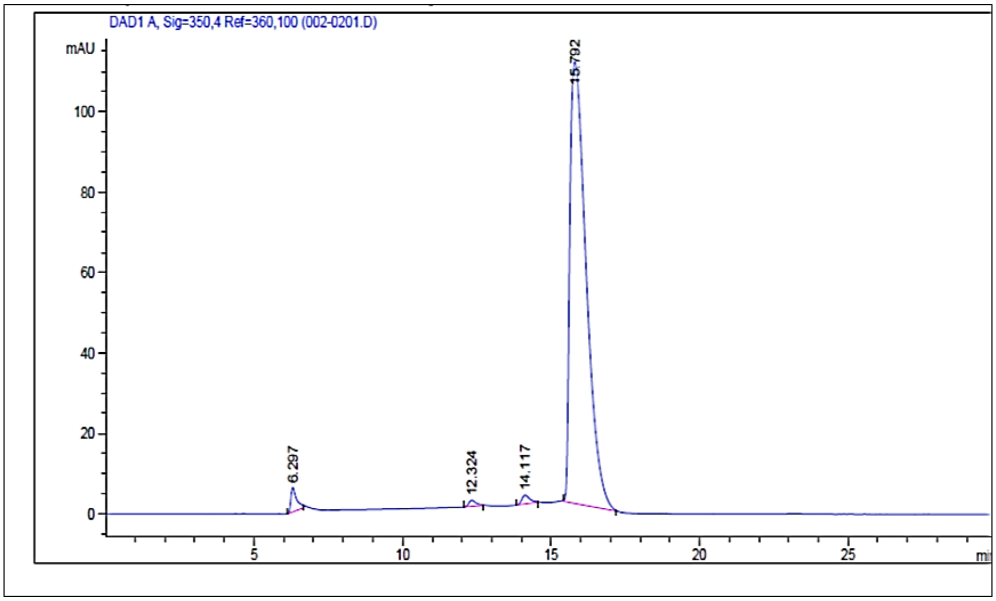


**Supplementary Figure 2:** The representative HPLC chromatogram of TAF, EVG, and DOX (DH) with retention times of ~5.35, 8.47, and 5.13 min, respectively**.**

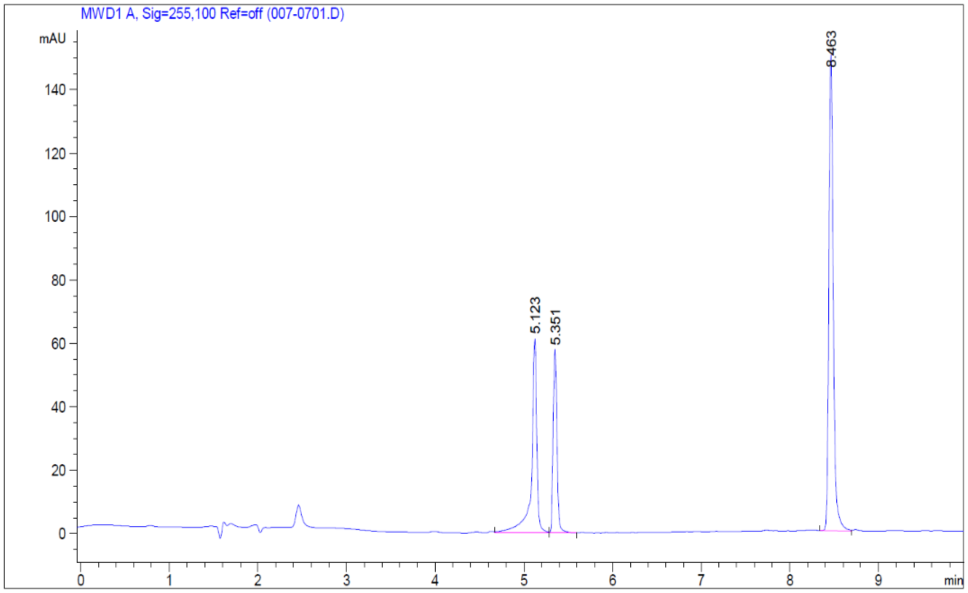
